# The bed nucleus of the stria terminalis mediates the expression of proactive defensive behavior

**DOI:** 10.1101/2022.08.16.504160

**Authors:** Diana P. Guerra, Wei Wang, Karienn A. de Souza, Justin M. Moscarello

**Affiliations:** Department of Psychological & Brain Sciences, Texas A&M University, College Station, TX, United States; Department of Neuroscience, Baylor College of Medicine, Houston, TX, United States; Department of Neuroscience and Experimental Therapeutics, Texas A&M Health Science Center, Bryan, TX, United States; Texas A&M Institute for Neuroscience (TAMIN), Texas A&M University, College Station, TX, United States

## Abstract

The bed nucleus of the stria terminalis (BNST) is a forebrain region implicated in aversive responses to uncertain threat. Much of the work on the role of BNST in defensive behavior has used Pavlovian paradigms in which the subject reacts to aversive stimuli delivered in a pattern determined entirely by the experimenter. Here, we report that BNST also mediates proactive defensive responses in a task that allows subjects to prevent the delivery of an aversive outcome. In a standard two-way signaled active avoidance paradigm, male rats learned to shuttle during a tone to avoid shock. Our data demonstrate that chemogenetic inhibition (hM4Di) of BNST attenuates the expression of the avoidance response, whereas chemogenetic activation (hM3Dq) of BNST potentiates the response by extending the period of tone-evoked shuttling. This effect was specific to the BNST, as inactivation of the neighboring medial septum produced no effect on the expression of avoidance. These data support the novel conclusion that BNST mediates two-way avoidance behavior in male rats.

## Introduction

An avoidant style of coping is a unifying behavioral feature of many forms of pathological anxiety^1^. Although healthy proactive avoidance responses prevent harm and reduce contact with stressors^2,3^, avoidant behavior that occurs at a relatively low cost can easily become excessive, fostering the maladaptive sensitivity to false alarm prevalent in anxiety disorders^4,5^. Despite the crucial role of avoidance in both adaptive defense and pathological anxiety, its neural circuitry has not been elucidated.

Two-way signaled active avoidance (SAA) is an acquired form of proactive avoidance in rodents. SAA involves a response (two-way shuttling) that is triggered by a conditioned stimulus (CS, tone) and serves to prevent contact with an aversive unconditioned stimulus (US, shock). As the response is acquired over the course of SAA training, US frequency decreases substantially. Viewed from the perspective of threat imminence theory^6-9^, this change in US density transforms the shock from a certain/imminent threat during initial acquisition to a possible/distal threat once the subject reaches asymptotic levels of SAA expression. Indeed, the CS will evoke high-imminence defensive reactions early in SAA training (freezing, ultrasonic vocalization, conditioned suppression, and conditioned analgesia), but these responses reduce as the expression of avoidance becomes more robust^10-15^. Despite evidence for a shift away from the defensive mode evoked by imminent threat, no previous research has explored the role played by the neural substrates of possible/distal threat in SAA.

Prior work demonstrates that conditioned reactions to distal or uncertain threat are underpinned by the bed nucleus of the stria terminalis (BSNT)^16-20^. We therefore hypothesized that the BNST mediates the expression of SAA. To test this hypothesis, we used a chemogenetic approach to either inhibit or activate BNST neurons in male rats performing this behavior. Our results demonstrate that inhibition of BNST attenuates the expression of SAA, whereas activation of this region produces a period of elevated responding that continues beyond the CS. These data establish BNST as a crucial node in the circuitry underlying proactive avoidance.

## Materials & Methods

### Animals

Subjects were 58 adult, male Sprague Dawley rats weighing at least 300 g at the beginning of experimentation. Rats were singly housed and maintained on a 14:10 light/dark cycle with *ad libitum* access to food and water in the vivarium in the Department of Psychological and Brain Sciences at Texas A&M University. Subjects were handled by experimenters before behavioral training. All procedures were conducted during the light phase and with the approval of the Texas A&M IACUC.

### Surgery

Animals were anesthetized with a 1mL/kg dose of a 10:1 mixture of ketamine (100mg/mL) and xylazine (100mg/mL) and mounted to a stereotaxic frame. Following incision, bilateral microinjections were made in BNST (ML +/-1.2, AP −0.12 or −0.24, DV - 7.3 or −7.6) or a unilateral microinjection was made in MS (ML +/-2.0, AP +0.36, DV −7.3 with cannula angled at 15 degrees off vertical). All infusions occurred at a rate of 0.1*𝜇*L/min for a total of 0.3*𝜇*L/injection and were followed by 5 min for diffusion. Each infusion contained an adeno-associated virus (AAV) from Addgene, bearing the gene construct for either an inhibitory or excitatory DREADD (Designer Receptor Exclusively Activated by Designer Drugs) or GFP. The following AAVs, concentrated at 1.2×10^13^ GC/ml, were used: AAV5-hSyn-hM4Di-mCherry, AAV5-hSyn-hM3Dq-mCherry, or AAV5-hSyn-EGFP, depending on the experiment. Animals were allowed at least 28 days of recovery before slice physiology or behavioral training.

### Electrophysiological Recordings

Coronal brain slices (250 μm) containing the BNST were taken from rats expressing hM4Di, hM3Dq, or GFP in BNST neurons. Slices were positioned in a perfusion chamber attached to the fixed stage of an upright microscope (Olympus) and submerged in continuously flowing oxygenated Artificial cerebrospinal fluid (aCSF) (in mM: 125 NaCl, 4.5 KCl, 2 CaCl_2_, 1.25 NaH_2_PO_4_, 25 NaHCO_3_, 15 sucrose and 15 glucose) at 32°C. Neurons were viewed under a water-immersion lens (40×) and a CCD camera. A Multiclamp 700B amplifier with Clampex 10.6 software and Digidata1555A (Molecular Devices) were used for recordings. The patch pipettes with resistance 3-5 MΩ were pulled by the pipette puller (Sutter Instrument Co., Model P-97). To measure spontaneous action potential (sAP), cell-attached voltage-clamp recordings of GFP or mCherry-positive putative GABAergic neurons in BNST were performed. Pipette was filled with the K^+^-based intracellular solution (in mM: 123 potassium gluconate, 10 HEPES, 0.2 EGTA, 8 NaCl, 2 MgATP and 0.3 NaGTP, pH 7.3, 280 mOsm). To record spontaneous EPSCs (sEPSCs), Picrotoxin (100 μM) and D-APV (25 μM) were added into the standard aCSF, and the neuron was held at a potential of −70 mV. Pipette was filled with the Cs-based intracellular solution (in mM: 119 CsMeSO_4_, 8 TEA. Cl, 15 HEPES, 0.6 EGTA, 5 QX-314.Cl, 7 phosphocreatine, 4 MgATP, 0.3 Na_3_GTP, 0.1 leupeptin, pH 7.3, 280 mOsm).

### Two-way Signaled Active Avoidance (SAA) Behavior

#### Training

Our SAA paradigm has been described elsewhere^14^. Briefly, subjects were trained to avoid by shuttling in either direction across a divided chamber during presentation of an auditory CS (15-sec, 2-kHz, 70-db pure tone). This response caused the immediate inactivation of the CS and the omission of the US (0.5-sec, 0.7-mA scrambled footshock).

#### Poor Avoider Criterion

Between training and test, data from all subjects in all experiments were reviewed and a poor avoidance criterion was applied^10,14,21^. Animals averaging six or fewer responses during the final two sessions of training were determined to be poor avoiders. The brains of poor avoiders were visually examined for any obvious lesion, but the extent of viral expression was not characterized.

#### Test Under Training Conditions

This procedure involved two additional days of two-way SAA with all training parameters in place. The only distinction between training and test was that ip injection of either CNO or vehicle was delivered in a counterbalanced order prior to each test session.

#### Test Under Extinction Conditions

To create a uniform behavioral assay with common conditions, this test was comprised of a single session in which ten 15-sec CSs were presented, each separated by a 2-min ITI. Unlike training, each CS continued for the full 15 sec regardless of whether or not the subject shuttled. No USs were presented. All subjects received ip injection of CNO or vehicle (for poor avoiders only) prior to test.

### DREADD Agonist

Clozapine-N-oxide (CNO; 3mg/kg/mL) was dissolved in physiological saline with 10% DMSO was administered via intraperitoneal (ip) injection 20-25 min before test. Vehicle injection was physiological saline with 10% DMSO, also administered ip.

### Perfusion and Viral Expression

At the end of each behavioral experiment, animals were deeply anesthetized and transcradially perfused with 10% buffered Formalin. Brains were removed and sliced into 40-μm sections using a cryostat. Sections were mounted onto subbed slides and cover-slipped with Fluoromount (Sigma Aldrich) before examination under a fluorescent microscope (Olympus U-RFL-T microscope with an Olympus DP72 digital color camera and a Pior *OptiScan II* control system) to reveal either mCherry or GFP expression.

### Behavioral Analyses

Shuttling data were collected automatically by the Coulbourn GraphicState software used to automate the delivery of stimuli during SAA training and test. Digital videos of all test sessions were also recorded. Using Noldus Ethovision software, these recordings were analyzed offline for freezing and locomotor activity data.

### Figures

Graphs were made using GraphPad Prism. Viral expression images were made with maps from the Swanson atlas^22^ using Adobe Illustrator. Behavioral schematics were made using BioRender.com.

## Results

### Chemogenetic Modulation of Neural Activity in BNST

To illustrate the effects of inhibitory and excitatory DREADDs on the activity of BNST neurons, we expressed hM4Di, hM3Dq, or GFP in the BNST of male rats (procedure schematized in FIG 1A, top). Because it has been reported that a subset of neurons in BNST are spontaneously active^23^, slices were created for patch-clamp recordings of spontaneous neural activity in fluorescent cells (mCherry for hM4Di and hM3Dq, or GFP). Robust expression of mCherry (example image in FIG 1A, bottom) or GFP were evident in all slices. Baseline cell-attached recordings were made before exposing the slice to the DREADD agonist clozapine-N-oxide (CNO). Bath application of CNO in aCSF had no effect on spontaneous action potentials (sAPs) in BNST neurons expressing GFP only (FIG 1B, top). In contrast, CNO decreased the frequency of sAPs in hM4Di-expressing neurons (FIG 1B, middle) and increased the frequency of sAPs in hM3Dq-expressing neurons (FIG 1B, bottom). Similarly, CNO did not change excitatory postsynaptic currents (EPSCs) relative to baseline in GFP-expressing neurons (FIG 1C, top), though it did result in decreased frequency and amplitude of EPSCs in hM4Di neurons (FIG 1C, middle) as well as increased frequency and amplitude of EPSCs in hM3Dq neurons (FIG 1C, bottom). Thus, CNO altered the activity of hM4Di- and hM3Dq-expressing BNST, and these effects were absent in GFP-expressing neurons.

**FIGURE 1.**
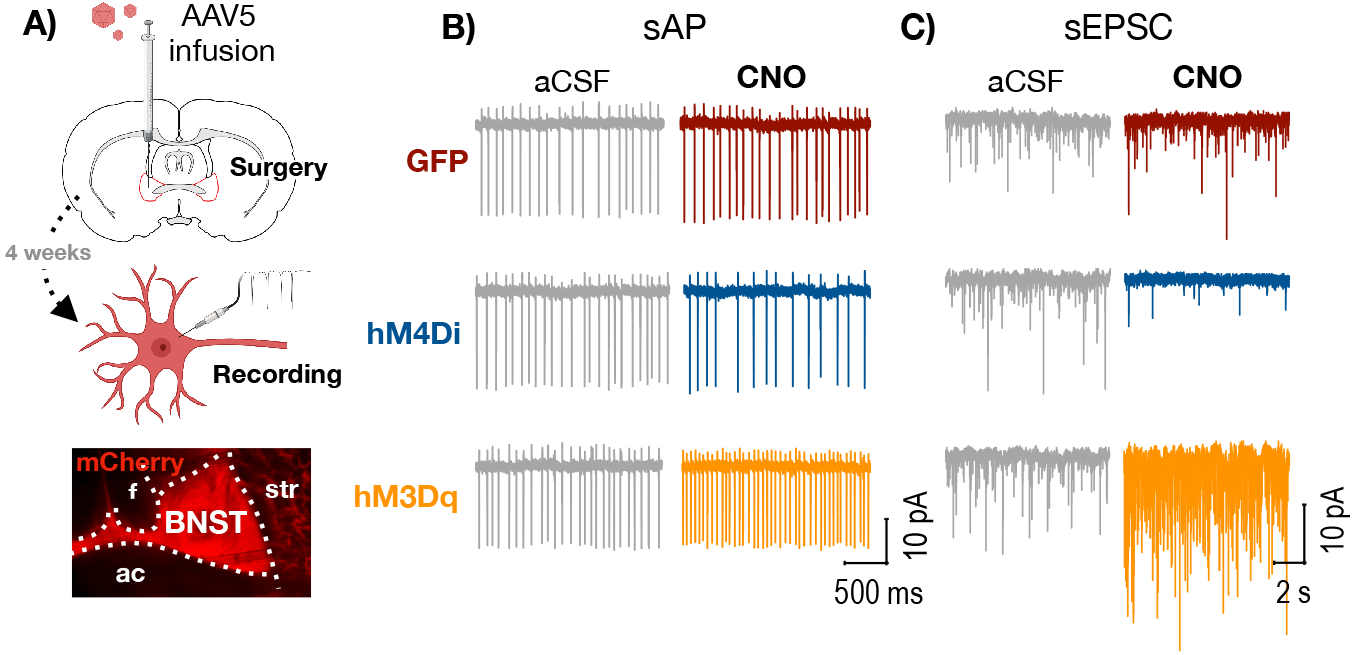
**A)** Top: subjects received intra-BNST infusions of AAV bearing the gene construct for either hM4Di, hM3Dq, or GFP and allowed to recover before recordings of spontaneous neural activity were conducted in slice. Bottom: representative image of mCherry expression in BNST (ac = anterior commissure, f = fornix, str = striatum). **B)** Representative cell-attached recordings of spontaneous action potentials (sAPs) in BNST neurons expressing GFP (top), hM4Di (middle), or hM3Dq (bottom). Bath application of 10μm of CNO had no effect on GFP-expressing neurons, but caused a decrease in sAPs in hM4Di-expresing neurons and an increase in hM3Dq-expressing neurons. **C)** Representative whole-cell recordings of spontaneous excitatory synaptic currents (sEPSCs) in BNST neurons expressing GFP (top), hM4Di (middle), or hM3Dq (bottom). Bath application of 10μm of CNO had no effect on GFP-expressing neurons, but caused a decrease in frequency and amplitude of sEPSCs in hM4Di-expressing neurons and an increase in hM3Dq-expressing neurons.

### Chemogenetic Inhibition of BNST Attenuates the Expression of SAA

To determine whether BNST is necessary for the expression of SAA, we expressed the inhibitory hM4Di DREADD or GFP in the BNST of male rats (FIG 2B depicts viral expression) that received four days of SAA training. Following training but before test, two rats were determined to be poor avoiders and were removed from our analysis, leaving us with the following groups: hM4Di (n=9) and GFP (n=8). These groups then underwent testing under training conditions in which two normal SAA sessions were preceded by either CNO or vehicle given in a counterbalanced order (design schematized in FIG 3A).

**FIGURE 2.**
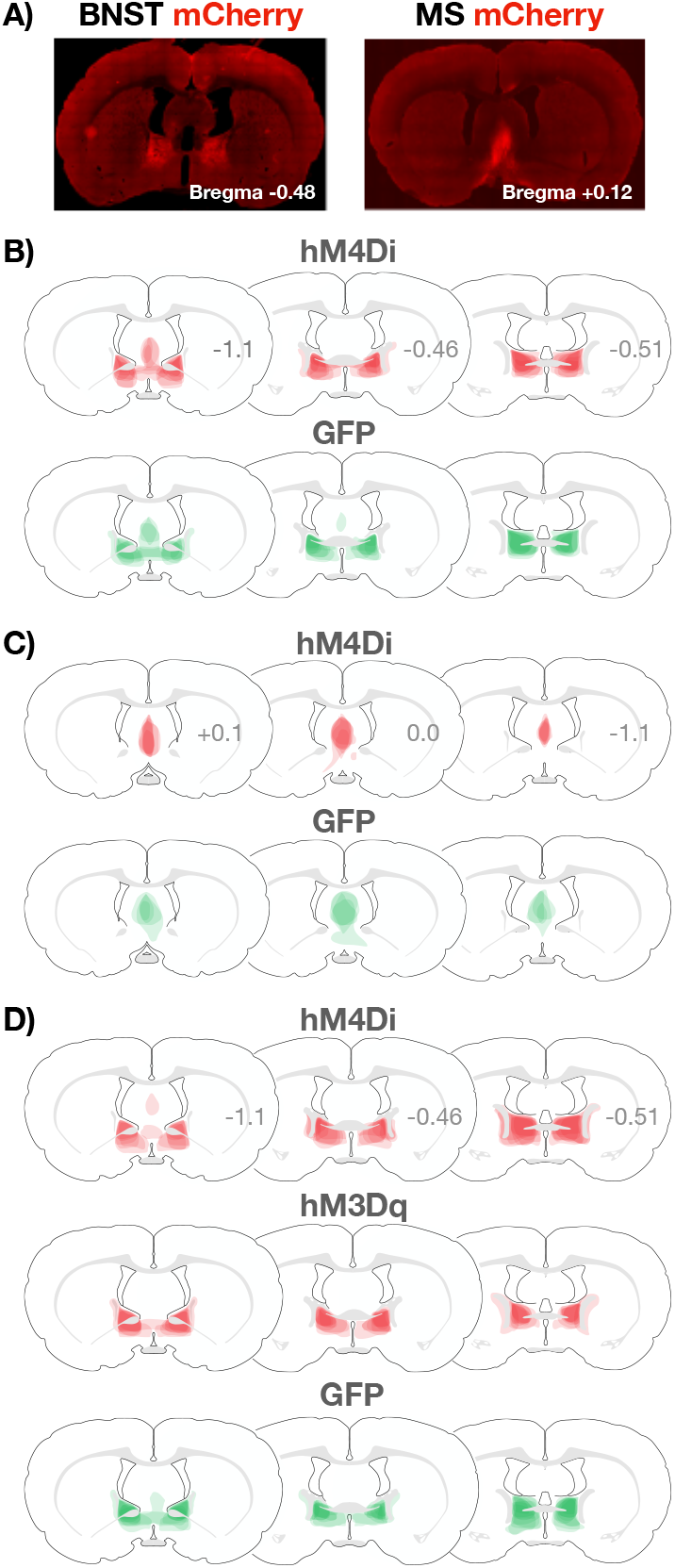
**A)** Representative images of mCherry expression in the BNST (left) and MS (right). **B)** Extent of viral expression in hM4Di (mCherry, top) and GFP (bottom) subjects from the experiment in which BNST was inactivated with CNO during a test conducted under training conditions (see FIG 3). **C)** Extent of viral expression in hM4Di (mCherry, top) and GFP (bottom) subjects from the experiment in which MS was inactivated with CNO during a test conducted under training conditions (see FIG 4). **D)** Extent of viral expression in hM4Di (mCherry, top), hM3Dq (mCherry, middle), and GFP (bottom) subjects from the experiment in which BNST was activated or inactivated with CNO during a test conducted under extinction conditions (see FIG 5).

**FIGURE 3.**
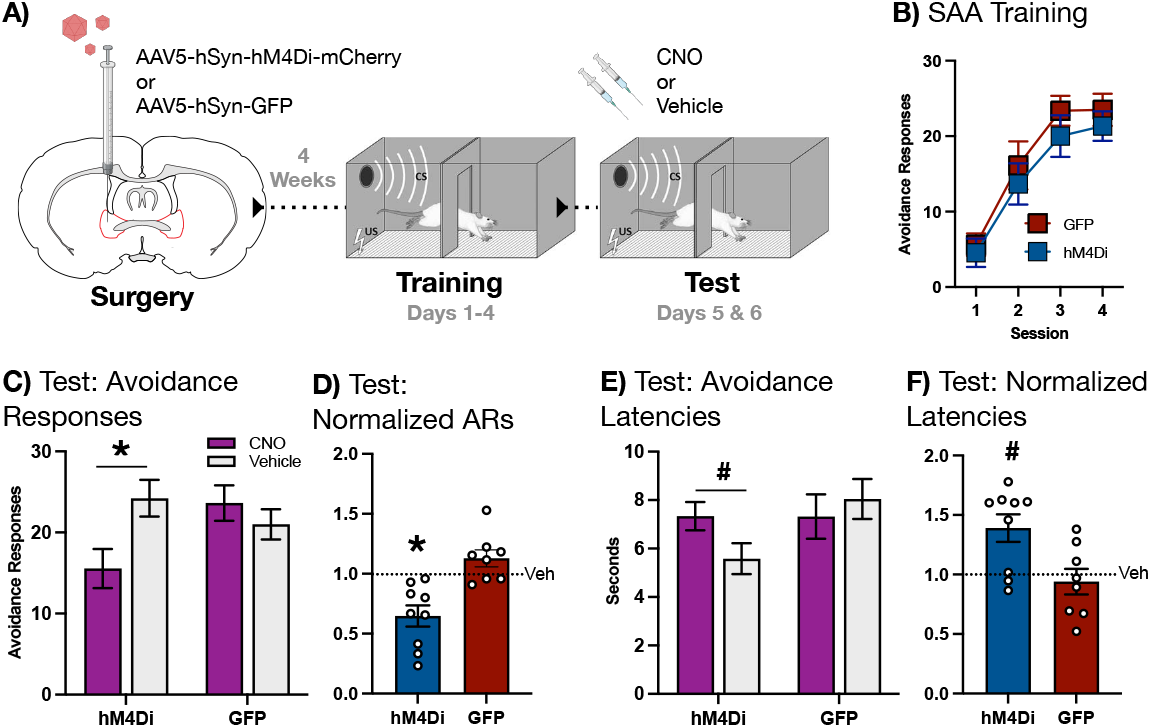
**A)** Subjects received intra-BNST infusions of AAV bearing the gene construct for either the hM4Di DREADD or GFP and allowed to recover. Following recovery, subjects underwent 4 days SAA training prior to a pair of tests conducted under training conditions, preceded by either CNO or vehicle. **B)** SAA training data. **C)** Avoidance responses at test: CNO significantly attenuated expression of the two-way avoidance response in hM4di-but not GFP-expressing subjects (**\***signifies p<0.01). **D)** Avoidance responses at test normalized to vehicle: CNO decreased the expression of avoidance in the hM4Di group relative to the GFP group (**\***signifies p<0.01). **E)** Avoidance latencies at test: CNO significantly increased the latency to avoid in hM4Di-but not GFP-expressing subjects (^**#**^ signifies p<0.05). **F)** Avoidance latencies at test normalized to vehicle: CNO significantly increased avoidance latency in the hM4Di group relative to the GFP group (^**#**^ signifies p<0.05). All graphs depict mean (±SEM).

Both hM4Di and GFP groups acquired SAA (FIG 3B). At test, CNO attenuated the expression of the avoidance responses in subjects expressing hM4Di, but not GFP (FIG 3C). A two-way, mixed-design ANOVA with a within-subjects factor of Drug (CNO or vehicle) and a between-subjects factor of Group (hM4Di or GFP) revealed a significant Drug X Group interaction [F(1,15)=15.95, p=0.0012]. Fisher’s LSD *post-hoc* tests further revealed that this interaction was driven by a significant reduction in avoidance responses when the hM4Di group received CNO relative to vehicle (p=0.00045). No such reduction was observed in the GFP group. To further examine the effects of BNST inactivation on avoidance, we normalized behavior during test by dividing each subject’s avoidance responses performed following CNO by avoidance responses performed following vehicle (FIG 3D). Two-tailed *t*-test conducted on these data confirmed a significant decrease in normalized avoidance responses in hM4Di relative to GFP subjects [*t*(15)=4.189, p=0.00079]. Thus, chemogenetic inhibition of BNST attenuated the expression of SAA.

To assess the residual avoidance responses that occurred despite the influence of BNST inactivation, we measured the latency to avoid during test. Administration of CNO increased avoidance latencies in subjects expressing hM4Di but not GFP (FIG 3E). A two-way, mixed-design ANOVA with a within-subjects factor of Drug (CNO or vehicle) and a between-subjects factor of Group (hM4Di or GFP) revealed a significant Drug X Group interaction [F(1,15)=5.529, p=0.033]. Fisher’s LSD *post-hoc* test further revealed that this interaction was driven by a significant increase in avoidance latency when hM4Di-expressing subjects received CNO relative to vehicle (p=0.028). To further explore the influence of BNST inactivation on the latency to avoid, we normalized each subject’s latencies during test (FIG 3F), as described above. Two-tailed *t*-test performed on these data revealed a significant increase in hM4Di subjects relative to GFP subjects [*t*(15)=2.831, p=0.013]. Thus, BNST inactivation reduced the number of avoidance responses at test and increased the latency to perform the residual responses that persisted in BNST-inactivated rats.

### Chemogenetic Inhibition of the Medial Septum Has No Effect on the Expression of SAA

Several subjects in the previous experiment showed mCherry-expression in the medial septum (MS) (FIG 2B). To control for the effect of hM4Di inactivation of MS on avoidance, we performed an experiment in which we explicitly tested the role of MS in SAA. The design of this experiment was identical to the previous, with the exception that hM4Di (n=6) or GFP (n=4) were expressed in MS (FIG 2C depicts viral expression). Subjects received four sessions of SAA training prior to a pair of tests under training conditions preceded in a counterbalanced order by CNO or vehicle (design schematized in FIG 4A). All subjects acquired SAA (FIG 4B). However, two-way mixed-design ANOVA revealed that CNO had no effect on the expression of the avoidance response (FIG 4C) or on avoidance latencies (FIG 4D).

**FIGURE 4.**
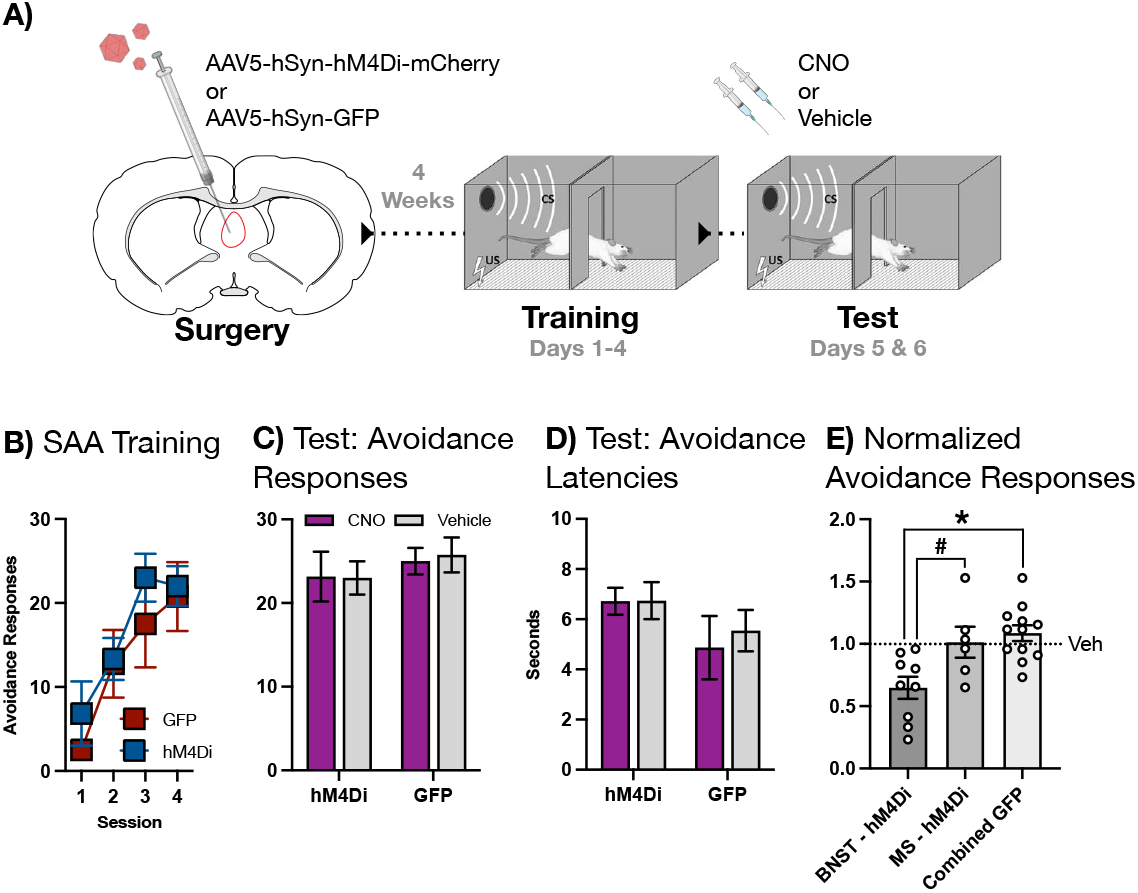
**A)** Subjects received intra-MS infusions of AAV bearing the gene construct for either the hM4Di DREADD or GFP and allowed to recover. Following recovery, subjects underwent 4 days SAA training prior to a pair of tests under training conditions, proceeded by either CNO or vehicle. **B)** SAA training data. **C)** Avoidance responses at test: CNO had no effect on the expression of the two-way avoidance response. **D)** Avoidance latencies at test: CNO had no effect on the latency to avoid. **E)** Avoidance responses normalized to vehicle in subjects expressing hM4Di in either BNST or MS compared to a combined BNST/MS GFP group. CNO decreased the expression of avoidance in the BNST-hM4Di group relative to the MS-hM4Di group (^**#**^ signifies p<0.05) and the combined GFP group (**\*** signifies p<0.01). All graphs depict mean (±SEM).

To firmly establish that our behavioral effects are BNST-specific, normalized avoidance scores for each subject from this and the previous experiment were subjected to a direct statistical comparison. This analysis revealed that CNO selectively decreased the expression of the avoidance response in the BNST-hM4Di group (FIG 4E). A one-way ANOVA comparing BNST-hM4Di, MS-hM4Di, and a combined GFP group revealed a significant main effect [F(2,24)=8.192, p=0.0019]. Fisher’s LSD *post-hoc* tests further revealed that this effect was driven by a significant decrease in normalized avoidance in the BNST-hM4Di group relative to both the MS-hM4Di (p=0.011) and the combined GFP group (p=0.00063). No significant difference was observed between MS-hM4Di and GFP groups. Thus, we conclude that the effect of CNO on SAA in our hM4Di-expressing subjects was specific to BNST.

### Chemogenetic Activation and Inhibition of BNST Have Opposite Effects on the Expression of SAA

Because changes in the expression of the avoidance response alter the pattern of tones and shocks delivered to subjects in the SAA paradigm, we explored the role of the BNST in a test conducted under extinction conditions, in which the delivery of stimuli is uniform across groups. This experimental setup was used to both validate our previous results with chemogenetic inhibition (hM4Di) and also to explore whether chemogenetic activation (hM3Dq) would potentiate the response.

We expressed either hM4Di, hM3Dq, or GFP in the BNST (FIG 2D depicts the extent of viral expression in all subjects). Subjects then received 6 days of SAA training, after which the poor avoider criterion was applied. There was a sufficient number of poor avoiders to create an independent group instead of disqualifying these subjects from testing. Thus, on the day after SAA training ended, the following groups underwent testing under extinction conditions: hM4Di (n=8), hM3Dq (n=8), GFP (n=9), and poor avoiders (n=6). All subjects received CNO prior to test, with the exception of the poor avoider group, which received vehicle (design schematized in FIG 5A).

**FIGURE 5.**
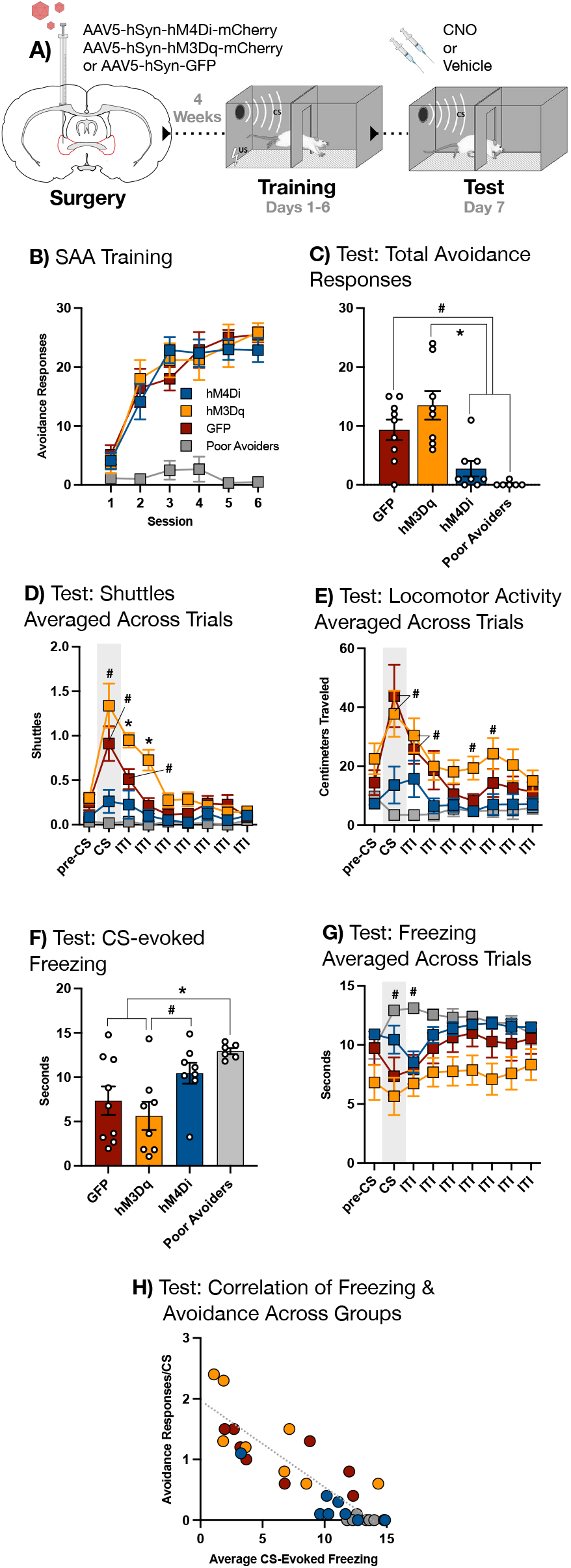
**A)** Subjects received intra-BNST infusions of AAV bearing the gene construct for either the hM4Di DREADD, the hM3Dq DREADD, or GFP and allowed to recover. Following recovery, subjects underwent 6 days SAA training prior to a single test conducted under extinction conditions (10 CSs, no USs), preceded by either CNO or vehicle (for poor avoiders only). **B)** SAA training data. **C)** Total avoidance responses (shuttles during the CS) at test: GFP controls produced more avoidance responses relative to both hM4Di subjects and poor avoiders (^**#**^ signifies p<0.05); hM3Dq subjects also produced more avoidance responses relative to both hM4Di subjects and poor avoiders (**\*** signifies p<0.01). **D)** Shuttles averaged across all 10 trials at test (x-axis divided into 15-sec epochs): CS presentation elevated two-way shuttling in hM3Dq subjects relative to their pre-CS baseline (^**#**^ signifies p<0.05), and this trend persisted into the first two epochs of the inter-trial interval (ITI) (**\*** signifies p<0.01); CS presentation also elevated shuttling in GFP controls, and this trend persisted into the first epoch of the ITI only (^**#**^ signifies p<0.05). **E)** Locomotor activity (centimeters traveled) averaged across trials at test: CS presentation increased locomotion in hM3Dq and GFP subjects relative to poor avoiders only, and this trend persisted into the first epoch of the inter-trial interval (ITI) (^**#**^ signifies p<0.05); locomotor activity was elevated in the hM3Dq group relative to both the hM4Di group and the poor avoiders during the 4^th^ epoch of the ITI and relative to the poor avoiders only during the 5^th^ epoch of the ITI (^**#**^ signifies p<0.05). **F)** Average freezing during the CS: poor avoiders froze more than GFP and hM3Dq subjects (**\*** signifies p<0.01); hM4Di subjects froze more than hM3Dq subjects (^**#**^ signifies p<0.05). **G)** Freezing averaged across trials at test: CS presentation increased freezing in poor avoiders relative to their pre-CS baseline, and this trend continued into the first epoch of the ITI (^**#**^ signifies p<0.05). **H)** Correlation of avoidance responses per CS and average freezing during the CS for all subjects (r= ^**-**^0.87, p<0.01). Other than the correlation scatter plot, all graphs depict mean (±SEM).

All groups showed normal avoidance during training, with the exception of poor avoiders (FIG 5B). Of the data generated at test, we first examined the effect of our chemogenetic manipulations on the expression of the avoidance response (FIG 5C). One-way ANOVA conducted on total avoidance responses revealed a main effect for group [F(3,27)=11.30, p=0.00005]. Fisher’s LSD *post-hoc* tests further revealed that this effect was driven by a significant decrease in avoidance responses in both the hM4Di and poor avoider groups relative to the GFP group (p=0.010 and p=0.0014, respectively). Fisher’s LSD *post-hoc* tests also demonstrated a significantly elevated response level in the hM3Dq group relative to the hM4Di and poor avoider groups (p=0.00015 and p=0.00003, respectively), but not GFP controls. These data replicate and extend the above finding that BNST inhibition attenuates the expression of the avoidance response.

To perform a more granular analysis of responses during test, we divided each trial into nine 15 sec epochs, starting with a pre-CS baseline period and continuing through the CS and inter-trial interval (ITI). The number of two-way shuttles (transitions from one side of the chamber to the other) occurring in each epoch was averaged across trials to create a representative picture of behavior across the test session (FIG 5D). These data were subjected to a two-way, mixed-design ANOVA with a between-subjects factor of Group (GFP, hM4Di, hM3Dq, or poor avoider) and a within-subjects factor of Time (epoch), which revealed a significant Group X Time interaction [F(24,208)=7.730, p<0.00001]. To interrogate the source of this effect, we used a Dunnett’s multiple comparisons test to compare the average number of shuttles during each group’s pre-CS baseline to every subsequent epoch. This analysis revealed that CS presentation significantly elevated two-way shuttling relative to the pre-CS baseline in both hM3Dq (p=0.016) and GFP (p=0.031) groups. Interestingly, hM3Dq subjects showed increased shuttling through the first and second post-CS epochs of the ITI (p=0.00031 and p=0.0048, respectively) before falling back to pre-CS baseline levels. In contrast, ITI shuttling in the GFP group remained marginally elevated above the pre-CS baseline during the first post-CS epoch only (p=0.0497). Notably, neither hM4Di nor poor avoider groups showed any significant deviation from their pre-CS baseline levels of two-way shuttling. Thus, the CS increased the expression of shuttling in animals receiving chemogenetic (hM3Dq) activation of BNST as well as in GFP control animals, and though this effect endured beyond the CS to some extent in both groups, it did so for longer in the case of BNST activation. In contrast, chemogenetic (hM4Di) inhibition of BNST eliminated the effects of the CS on expression of two-way shuttling, producing a pattern of behavior highly similar to poor avoiders, despite the fact the hM4Di subjects showed robust avoidance during training.

Next, we analyzed locomotor activity (distance traveled in cm) averaged across the same time bins as above (FIG 5E). This allowed us to assess whether changes in shuttling might be due to general enhancements or impairments in locomotion caused by our manipulations of BNST. Locomotor activity data were subjected to a two-way, mixed-design ANOVA with a between-subjects factor of Group and a within-subjects factor of Time, as above. This analysis revealed a significant Group X Time interaction [F(24,208)=3.115, p<0.00001], which we parsed using a Dunnett’s multiple comparisons test in which distance traveled during each group’s pre-CS baseline was contrasted with every subsequent time bin, identical to the *post-hoc* analysis described above. Intriguingly, this analysis uncovered no statistically significant deviations from the pre-CS baseline in any group. To uncover the source of the interaction revealed by the ANOVA, we then conducted a Tukey’s multiple comparisons test to examine how locomotor activity differed between groups within each epoch. Notably, there were no statistically significant differences between any group during the pre-CS baseline. During the CS, significant differences in distance traveled were observed between poor avoiders and the hM3Dq and GFP groups (p=0.013 and p=0.026, respectively). The difference between poor avoiders and hM3Dq subjects was also present in the 1^st^ and 2^nd^ ITI epochs (p=0.0089 and p=0.043, respectively). In contrast, the difference between poor avoiders and GFP controls and was only significant in the 1^st^ ITI epoch following the CS (p=0.0099). In addition, significant differences between poor avoider and hM3Dq groups remerged during the 4^th^ and 5^th^ ITI following the CS (p=0.032 and p=0.036, respectively). The only significant difference that did not involve the poor avoiders was between hM3Dq and hM4Di groups during the 4^th^ ITI epoch after the CS (p=0.032). Thus, the differences revealed in this analysis seem to be driven by reduced locomotion in poor avoiders more than any other factor (though, we note again that the pre-CS baseline was the same for all groups). We conclude that the effects of BNST activation and inhibition on shuttling cannot be attributed to alterations in locomotion.

We also measured CS-evoked freezing at test (FIG 5F). Time spent freezing was averaged over ten CSs for each subject and then analyzed with a one-way ANOVA that revealed a significant main effect for Group [F(3,27)=4.917, p=0.0075]. Fisher’s LSD *post-hoc* tests further revealed that the hM4Di group froze more than the hM3Dq group (p=0.0199), though freezing in hM4Di subjects was not significantly different than GFP controls or poor avoiders. In addition, Fisher’s LSD *post-hoc* tests revealed that poor avoiders froze significantly more than both hM3Dq subjects (p=0.0018) and GFP controls (p=0.011), consistent with the high level of freezing previously observed in subjects that fail to express the avoidance response^10,14^. Neither chemogenetic activation nor inhibition altered freezing during the CS relative to GFP controls, suggesting that changes in the avoidance response caused by these manipulations cannot be accounted for by changes in freezing.

Next, we again broke each trial into 15-sec epochs starting with a pre-CS baseline and averaging the time spent freezing during each consecutive epoch across trials (FIG 5G). These data were analyzed using a two-way, mixed-design ANOVA with a between-subjects factor of Group (GFP, hM4Di, hM3Dq, and poor avoider) and a within-subjects factor of Time (epoch), which revealed a significant Group X Time interaction [F(24,216)=4.127, p<0.00001]. We then interrogated this effect using a Dunnett’s multiple comparisons test to compare time spent freezing during each group’s pre-CS baseline to freezing in all subsequent epochs. The only significant differences that this analysis revealed were in the poor avoider group, which froze more relative to the pre-CS baseline during both the CS and the 1^st^ epoch of the ITI following the CS (p=0.043 and 0.048, respectively). Freezing did not differ significantly from pre-CS baseline in any other group. The overall flatness of freezing across the trial may be due to the fact that the training environment acquires an aversive association strong enough to produce substantial levels of contextual freezing. However, this relatively invariant pattern of trial-wide freezing did not interfere with group-level differences in shuttling or avoidance.

Because prior research shows that manipulations which attenuate avoidance also tend to produce concurrent increases in CS-evoked freezing^15,24^, we analyzed the correlation between avoidance responses/CS and freezing/CS in all subjects (FIG 5H). Pearson’s r-coefficient analysis revealed a strong negative correlation between the two behaviors [r= ^**–**^0.8657, p<0.00001]. Thus, at the level of individual subjects, there was an inverse correlation between freezing and avoidance, even though group-level differences in freezing cannot account for the effects of our chemogenetic manipulations on the avoidance response.

Overall, these results confirm our first experiment, which demonstrated that inhibition of BNST impaired performance of the avoidance response, and extend that result by revealing that activation of BNST can produces a pattern of elevated responses that persists beyond the CS.

## Discussion

The data presented here are the first demonstration that the bed nucleus of the stria terminalis (BNST) is not only necessary for the expression of two-way signaled active avoidance (SAA) but also sufficient to enhance the output of this response. Previous work on the role of BNST in aversive associative learning paradigms has focused on respondent or reactive behaviors, most commonly freezing^18,19,25-34^, but also conditioned suppression^35^, fear-potentiated startle^36^, and flight^37^. Evidence for a role of BNST in SAA augments our understanding of this region by establishing for the first time its role in proactive defensive behavior. These heterogeneous behavioral functions are reflected in the anatomical heterogeneity of the BNST, which a collection of small, complexly interconnected sub-nuclei comprised of multiple cell types^38-41^. Distinct populations within specific BNST sub-nuclei have already been shown to underpin specific defensive responses^42-45^, suggesting that contrasting proactive and reactive behavioral functions may indeed be supported by dissociable mechanisms within BNST, similar to what meticulous circuit-dissection work has revealed of a related structure, the central amygdala^46-49^. Future research will parse the circuits underlying heterogenous repertoire of BNST-dependent defensive behavior.

Recent empirical^50^ and theoretical^51^ work argues that two-way avoidance should be categorized as a pre-encounter defensive behavior within the framework of threat imminence theory, an important conceptual tool for understanding conditioned aversion from the perspective of predator/prey dynamics in natural settings^6^. A threat imminence approach to SAA draws on ecological models of predatory defense that place proactive/preventative avoidance measures at a time point in predator/prey interactions occurring before the predator has been directly encountered, when the threat of attack is relatively distant^7,8,52^. Characterizing two-way avoidance as a component of the pre-encounter defensive mode fits comfortably with an important theme of previous research on BNST, which is that BNST-dependent conditioned defensive responses tend to be evoked by threats that are distal, ambiguous, or otherwise difficult to predict (e.g.^18,19,36,37^). As a substrate of distal threat processing, the BSNT may be recruited to SAA by the increasing psychological distance of the US that occurs as the rising frequency of the avoidance response decreases the frequency of shock, transforming a threat that is certain during early trials into one that is merely possible once the subject achieves stable levels of avoidance expression.

In summary, we demonstrate that chemogenetic inhibition or activation of BNST neurons can attenuate or potentiate the expression of SAA, respectively. These results are the first demonstration of a role for BNST in proactive avoidance behavior. Ongoing research will examine the relationship between threat imminence, avoidance, and the function of BNST.

## Acknowledgements

This work was supported by NIH grant R21MH126327 (JMM) and BBRF NARSAD Young Investigator Award 25196 (JMM). We thank Dr. Jun Wang for providing the CNO used for the slice physiology experiment.

## Author Contributions

Diana Guerra assisted in the design of the behavioral experiments, executed those experiments, assisted in the analysis of behavioral data, performed the relevant histology, and contributed to writing the manuscript. Wei Wang and Karienn Souza designed and executed the physiological experiment and edited the manuscript. Justin Moscarello designed the behavioral experiments and supervised their execution, analyzed behavioral data, and wrote the manuscript.

